# A *nox2* / *cybb* zebrafish mutant with defective neutrophil ROS production displays normal initial neutrophil recruitment to sterile tail injuries

**DOI:** 10.1101/2024.02.02.578645

**Authors:** Abdulsalam I. Isiaku, Zuobing Zhang, Vahid Pazhakh, Graham J. Lieschke

## Abstract

Reactive oxygen species (ROS) are important effectors and modifiers of the acute inflammatory response, recruiting neutrophils to sites of tissue injury. In turn, neutrophils are both consumers and producers of reactive oxygen species. Stimulated neutrophils generate reactive oxygen species in an oxidative burst through the activity of a multimeric phagocytic NADPH oxidase complex. Mutations in the *NOX2/CYBB* (previously gp91^phox^) NADPH oxidase subunit are the commonest cause of chronic granulomatous disease, a disease characterized by infection susceptibility and an inflammatory phenotype. To model chronic granulomatous disease, we made a *nox2/cybb* zebrafish (*Danio rerio*) mutant and demonstrated it to have severely impaired neutrophil ROS production. Reduced early survival of *nox2* mutant embryos indicated an essential requirement for *nox2* during early development. In *nox2/cybb* zebrafish mutants, the dynamics of initial neutrophil recruitment to both mild and severe surgical tailfin wounds was normal, suggesting that excessive neutrophil recruitment at the initiation of inflammation is not the primary cause of the “sterile” inflammatory phenotype of chronic granulomatous disease patients. This *nox2* zebrafish mutant adds to existing in vivo models for studying neutrophil reactive oxygen species function in development and disease.

## Introduction

Neutrophil granulocytes are major initial responders to infection and inflammation. Many neutrophil functions are regulated by intracellularly-generated reactive oxygen species (ROS). These include chemotaxis, phagocytosis, and neutrophil extracellular trap release. ROS production by neutrophils is generated by an active process involving the phagocyte NADPH oxidase (PHOX) enzyme. Neutrophil PHOX is a multi-subunit enzyme with cytosolic (NCF1/p47^phox^, NCF2/p67^phox^, NCF4/p40^phox^) and transmembrane (CYBA/p22^phox^, NOX2/CYBB/gp91^phox^) subunit proteins (Panday et al., 2015; Quinn, 2013; Thomas et al., 2017). CYBC1/EROS chaperones gp91^phox^ and p22^phox^ heterodimerization in the endoplasmic reticulum, but is not part of the NADPH oxidase complex itself (Randzavola et al., 2022).

The importance of PHOX-derived ROS is highlighted by chronic granulomatous disease (CGD), a primary immune deficiency characterized by genetic mutation in one of the neutrophil PHOX subunit proteins, severe recurrent infections and/or excessive inflammation (Dinauer, 2019). NOX2/CYBB is the most highly expressed PHOX subunit protein in neutrophils, and *NOX2/CYBB* mutations account for ∼65% of human CGD (Dinauer, 2019). Although the vulnerability of CGD patients to infections is directly linked to a reduced level of neutrophil NOX2-derived ROS production, the exact mechanisms behind inflammatory manifestations of CGD are less clear (Marciano et al., 2018; Roos, 2016).

Zebrafish models have made important contributions to understanding the roles of neutrophil-derived ROS. Several studies have used zebrafish morphants targeting *nox2/cybb* or other neutrophil PHOX subunits, showing increased susceptibility to fungal and bacterial infections attributed to ROS deficiency (Brothers et al., 2011; Mesureur et al., 2017; Yang et al., 2012). This is consistent with the established view that ROS-dependent neutrophil antimicrobial effects are central to host defense against pathogens and demonstrates the conserved role of ROS in cell-autonomous innate immunity (Randow et al., 2013). Two zebrafish *nox2/cybb* mutant alleles have been previously described, but their effects on neutrophil ROS production and other neutrophil cellular functions have not been characterized (Terzi et al., 2021; Weaver et al., 2018).

After tissue injury, ROS levels represent a complex interplay between ROS generated by other tissues and ROS consumption and production by neutrophils. Tissue injury results in a burst of DUOX-dependent burst of hydrogen peroxide (H_2_O_2_) production by epithelial cells which acts as a neutrophil chemoattractant (Niethammer et al., 2009) which is sensed in neutrophils by oxidation of the cysteine C466 residue in Lyn, a Src family kinase (Niethammer et al., 2009; Yoo et al., 2011). Arriving neutrophils down-regulate this H_2_O_2_ by a myeloperoxidase-dependent mechanism (Pase et al., 2012). However, activation of arriving neutrophils stimulates their oxidative burst, adding neutrophil-generated ROS to the inflammatory response. Understanding this dynamic interplay of ROS production, consumption and decay requires tools that selectively modulate the various ROS contributions of different cells and enzymes.

We have generated and characterized a new *nox2/cybb* zebrafish mutant with neutrophils displaying profoundly impaired ROS production. We also investigated the effect of their *nox2/cybb*-dependent ROS deficiency on initial neutrophil recruitment to acute surgical wounds, but found that loss of *nox2*-derived ROS did not alter the numbers of neutrophils initially recruited to minor or severe acute wound injuries. This mutant is a new model of CGD and provides a tool for studying Nox2-dependent myeloid cell dysfunction characterizing CGD.

## Materials and Methods

### Animal ethics and zebrafish alleles

Zebrafish strain used were: wild type (WT) Tübingen (TU) (Max-Planck-Institut für Entwicklungsbiologie, Tübingen, Germany), *Tg(mpx:EGFP)*^*i114*^ (Renshaw et al., 2006), the compound transgenic reporter line *Tg(mpx:Kal4TA4)*^*gl28*^ (Okuda et al., 2015), *Tg(UAS-E1b:Eco:NfsB-mCherry)*^*c26*4^ (Davison et al., 2007) and *nox2/cybb*^*gl43*^ (hereafter called *nox2*^*-/-*^) carried on both reporter backgrounds (this report). Zebrafish experiments were conducted under protocols approved by Monash University Animal Ethics Committees (MARP-2015-094/14375 and 17270) and in accordance with Australian Code of Practice for the Care and Use of Animals for Scientific Purposes (NHMRC, 2013).

### Gene expression analysis

The publicly-available EMBL-EBI Elixir node Single Cell Expression Atlas data and tools were employed using the following datasets (www.ebi.ac.uk/gxa/sc/home, accessed analysis 12/Jan/2024): “Single-cell RNA-seq analysis of kidney marrow from six zebrafish transgenic lines that label specific blood cell types” n=245 (Tang et al., 2017) and “Single-cell RNA-Seq data of zebrafish blood cells data” n=1,354 cells (Athanasiadis et al., 2017) and “Single-cell RNA sequencing of the cut and uncut caudal fin of zebrafish larvae” n=2,860 cells (data provided without published reference).

### CRISPR-Cas9 mutagenesis

Targeted mutation of *nox2* (Ensembl ENSDARG00000056) was achieved by microinjecting dual guide RNAs (gRNAs) complexed with Cas9 into one-cell stage Tuebingen (TU) zebrafish embryos (Jao et al., 2013). The gRNAs targeted sequences adjacent to PAM sites in exons 3 and 4 of the *nox2* gene (5′-GCCCACGAGAGAGCATGCTG-3′ and 5′-CAAGCTGTCGAGCTGCAGTG-3′ respectively).

### PCR and RT-PCR

Genomic DNA was extracted using HotShot method (Meeker et al., 2007). PCR amplification of fragments spanning the gRNA target sites used primers: gRNA1 site 5′-GGTTGTAAATGTGATGCCGTAA-3′ and 5′-AATTTCGGATACAGCCCAAGTA-3′; gRNA2 site, 5′-TGCAATCATAATGAAAAGGGA-3′ and 5′-GCTGCAATTCTTAAATATCCGC-3′. PCRs used Phusion High-Fidelity DNA Polymerase (ThermoFisher Scientific) and a T100 thermal cycler (Bio-Rad).

RNA was extracted with TRIzol reagent. cDNA was synthesized using SuperScript™ VILO cDNA Synthesis kit (ThermoFisher Scientific). RT-PCR to detect *nox2* missplicing used primers: *nox2* (positioned in introns 2 and 3), 5′-GTATGGCTCGGGATCAATGTGT-3′ and 5′-GATCCTCGGAGAAACGAGAGC-3′; *ppial*, 5′-ACACTGAAACACGGAGGCAAAG-3′ and 5′-CATCCACAACCTTCCCGAACAC-3′ (van der Vaart et al., 2013; van Soest et al., 2011).

### Sanger sequencing

The PCR products representing genomic and cDNA sequences were purified using PCR purification kit (Promega) and subsequently sequenced using Sanger sequencing at Micromon Genomics, Monash University.

### Quantification of ROS

ROS production in whole embryos was measured as previously described (Goody et al., 2013). Briefly, whole embryos of known genotype were divided into phorbol myristate acetate (PMA) and non-PMA stimulated groups in a dark 96 well plate. The redox dye 2,7dichlorodihydrofluorescein diacetate (H2CFDA) was added. Fluorescence readings were taken at room temperature after 30-60 min, every 12 min for 180 min using a spectrophotometer (Infinite M200 PRO, TECAN).

A ROS assay using dihydrorhodamine 123 (DHR123) was employed, as used in CGD diagnosis (Emmendorffer et al., 1994; Vowells et al., 1996). Phorbol 12-myristate 13-diacetate (PMA) was used to stimulate ROS production in FACS-purified myeloid cells obtained from WT and *nox2*^-/-^ zebrafish whole kidney marrow (WKM) prepared from non-transgenic backgrounds devoid of reporter genes. Unstimulated WKM myeloid cells served as control. The stimulation index was calculated by expressing the paired PMA stimulated:unstimulated fluorescence intensity ratio as a percentage. For FACS-purification, single cell preparations were prepared from dissected whole kidneys by mechanical disruption and strained through a 35 μm filter into FACs tubes. Dead cells were excluded by DAPI staining. The myeloid cell gate was determined by forward and side scatter flow analysis using BD LSRFortessa X20 (Traver et al., 2003). Samples are WKM myeloid cells from single animals.

### Viability assay

Zebrafish embryos from three adult zebrafish crosses, *WT x WT, WT x nox2*^*-/-*^ and *nox2*^*-/-*^ x *nox2*^*-/-*^ were observed at 24 hours or daily for up to 5 days post-fertilization. The numbers of live and dead embryos were counted manually.

### Wound assay

Two to three-day post fertilization embryos were anesthetized using 160 mg/mL tricaine methanesulfonate (Sigma-Aldrich). Using a fine scalpel blade, two types of wound were created of different severity, excising either the tip of the tail fin alone, or the tip of the tail fin along with the distal tip of the notochord. Images of individual embryos were taken using MVX10 microscope fitted with Olympus DP72 camera and CellSens software (version 1.11). The number of neutrophils at the wound site (caudal vein loop to transection point) was manually counted from images of embryos as previously described (Isiaku et al., 2021; Miskolci et al., 2019). Neutrophil numbers were scored blinded, prior to genotyping of embryos.

### Statistical analysis

GraphPad Prism Version 8.3.1 was used for statistical analysis. A two-way ANOVA with Tukey’s multiple comparison or unpaired two-tailed t-test were used to test the difference in ROS production. Mann-Whitney was used in testing differences in ROS stimulation index. Log-rank (Mantel-Cox) and Fisher’s exact tests were employed to compare survival rates. A two-way ANOVA with Geisser-Greenhouse correction and Tukey’s multiple comparisons tests was used to assess the relationship of time and number of neutrophils between the three groups of different genotype-matched siblings. Equality of variance and sphericity were not assumed. P values <0.05 were considered significant.

## Results

### Expression of zebrafish *nox2/cybb*

Zebrafish *nox2/cybb* (hereafter *nox2*) expression was evaluated using publicly-available single cell RNAseq datasets with reference to expression of the neutrophil marker genes *mpx* and *lyz*, and macrophage marker genes *mpeg1.1* and *mfap4* (Fig S1). Zebrafish *nox2* was expressed primarily in both phagocyte types, as expected of a phagocyte NAPDH oxidase (Fig S1A). It had a narrower cellular spectrum of expression than that of its transmembrane partner *cyba*/p22^phox^ (Fig S1B), consistent with the known functional partnership of Cyba with other Nox proteins (Quinn, 2013). The cytosolic subunit *ncf1/p47*^*phox*^ of phagocytic NAPDH oxidase displayed a similar pattern of expression. Within the heterogenous population of cells in the zebrafish tail fin region (Fig S1B), *nox2* expression segregated to a myeloid cluster containing *mpx-, ly-z, mpeg1.1-* and *mfap4-expressing* cells, whereas *cyba* displayed expression not restricted to this myeloid cluster, and *nox1* expression localised to an alternative cluster characterized by expression of marker genes potentially indicating an epithelial-related identity (e.g. *cyt1*).

### A zebrafish *nox2* mis-splicing mutant

To generate a zebrafish *nox2* mutant, two multiplexed gRNAs were delivered targeting PAM sites in intron 2 adjacent to the exon 3 splice acceptor and exon 4 of *nox2* (Fig 1A). The gRNAs were designed intending to induce deletion of intervening sequences between the two PAM targets, but no stable mutants with large deletions were recovered. A majority of F1 animals recovered carried mutations at either one or both PAM targets.

**Figure 1.**
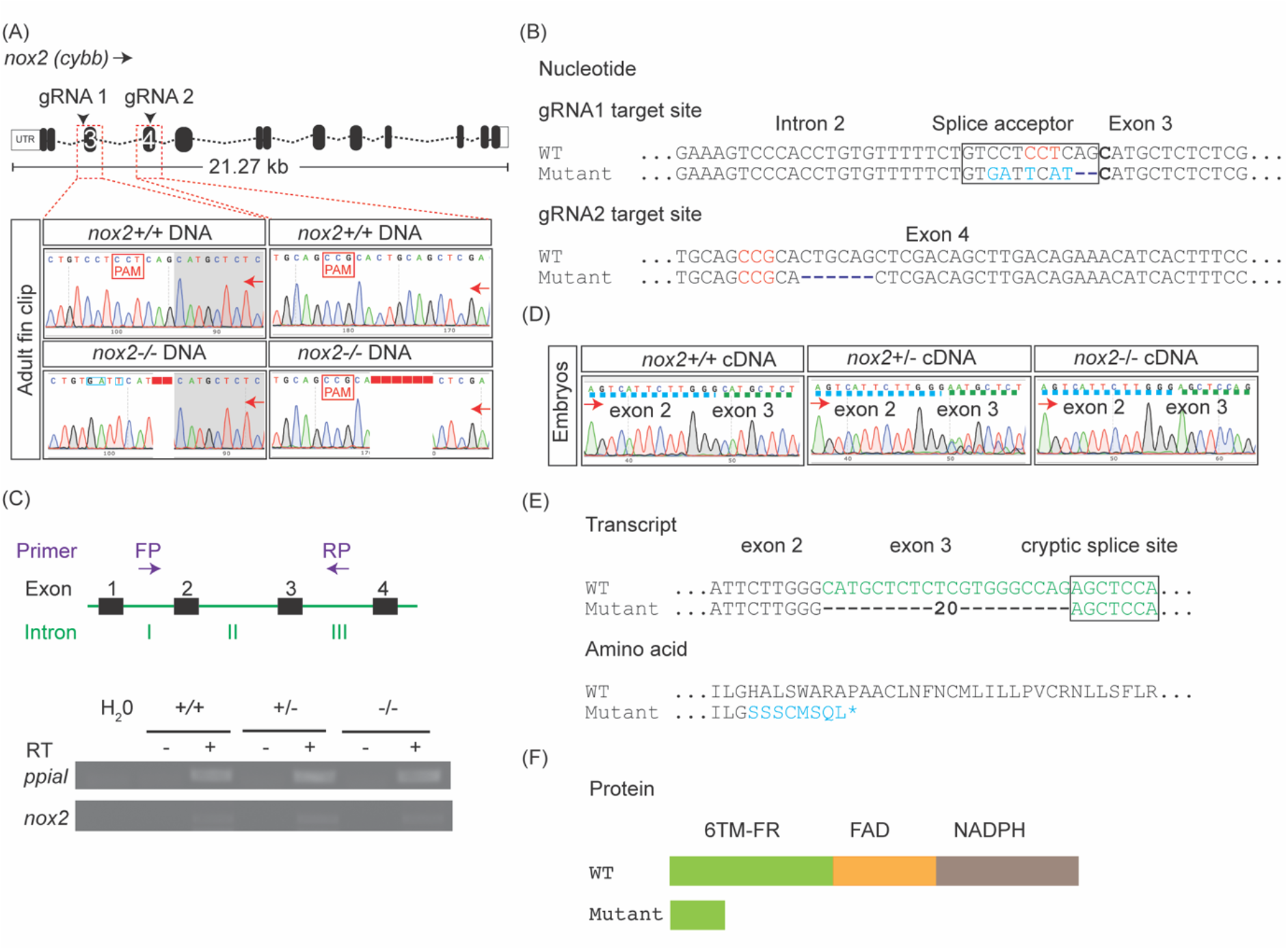
Genetic and molecular characterization of adult *nox2* zebrafish. (A)Sanger sequencing chromatogram of genomic DNA at gRNA 1 (intron 2/exon 3) and gRNA 2 (exon 4) target sites in adult wild type (upper panel) and *nox2* (lower panel) homozygous zebrafish. DNA sequence disruptions are around the protospacer adjacent motif (PAM) – NGG. (B)Interpretation of nucleotide sequences in A, at the gRNA 1 and gRNA 2 target sites (C)Schematic of PCR forward primer (FP) and reverse primer (RP) binding sites for amplifying cDNA transcripts (above). Gel image showing RT-PCR products from *nox2* wild type, heterozygous and homozygous zebrafish embryos (below) (D)Sanger chromatogram showing cDNA sequence traces of *nox2* wild type, heterozygous and homozygous zebrafish embryos from RT-PCR in C. Exons separated by coloured broken lines. (E)Interpretation of nucleotide sequence, and predicted amino acid sequence of wild type and *nox2* homozygous in D (F)Schematic of predicted protein domains in wild type and *nox2* zebrafish. *nox2*, NADPH oxidase 2; gRNA, guide RNA; PAM, protospacer adjacent motif; 6TM-FR, heme containing six transmembrane ferritin reductase domain; FAD, flavine adenine dinucleotide domain; NADPH, nicotinamide adenine dinucleotide phosphate domain; +/+, wild type (WT); +/-, heterozygous; -/-, homozygous; green font, exon 3 sequence; blue font, sequence mismatch; asterisk (*), premature stop; red arrows, direction of sequencing; FP, forward primer; RP, reverse primer; RT, reverse transcriptase.

A single mutant allele was selected for further studies. At the gRNA1 target site, there was a 2 base pair (bp) nucleotide deletion and 5 bp nucleotide substitution (Fig 1A, B). At the gRNA2 target site, there was a 6 bp deletion (Fig 1A, B). While the triplet deletion at the gRNA2 PAM did not alter the reading frame and was therefore of uncertain functional significance, the changes at the gRNA1 target site altered the exon 3 splice acceptor site, potentially leading to a transcript encoding a functionally defective protein.

To investigate whether mis-splicing occurred, RT-PCR was performed on cDNA from wild type, heterozygous and homozygous mutant *nox2* zebrafish embryos. Although electrophoresis of RT-PCR products provided no evidence of intron retention (Fig 1C), the products were sequenced to detect other forms of alternative splicing. This demonstrated that the mutation caused 20 bp of exon 3 sequence to be abnormally spliced out in the *nox2*^-/-^ zebrafish (Fig 1D, E). Although a mixture of normal and misspliced sequences were evident in heterozygous animals, in homozygous mutant animals, only the mis-spliced transcript was detected by Sanger sequencing. The mis-spliced sequence results in a frameshift leading to a premature stop codon, encoding a truncated Nox2 protein lacking the entire FAD and NADPH domains, with resultant loss of function. Hence, it was predicted that the homozygous mutants would be functionally Nox2-deficient.

### Impaired ROS production by *nox2* mutant neutrophils

ROS production was first assessed by PMA-stimulated whole embryos. In this assay, ROS production in WT embryos was completely PMA-dependent, precluding interference from EGFP reporter genes carried in the genetic background (Fig S2A). PMA stimulated ROS activity in all WT embryos, with variation between individual embryos. (Fig S2A). In the context of this variation, in this whole embryo assay, both *nox2*^*+/-*^ and *nox2*^*-/-*^ embryos displayed a range of PMA-stimulated ROS production that overlapped with that of WT embryos (Fig S2B).

We hypothesized that the loss of Nox2-dependent ROS production in *nox2*^*-/-*^ embryos would be predominantly in myeloid cells and would potentially be masked in whole embryo assays by widespread PMA stimulation of ROS production by other NADPH oxidases, including Nox1 (Kim et al., 2007), Nox3 (Ueno et al., 2005) and Nox5 (Jagnandan et al., 2007). ROS production was therefore assessed in myeloid cells specifically, using FACS-purified myeloid cells from adult whole kidney marrow (gating strategy shown in Fig S3) and assessing ROS production by a flow cytometric assay based on the oxidative spectral shift of DHR123 following PMA stimulation of the neutrophil oxidative burst. PMA-stimulated wild type myeloid cells showed robust induction of ROS activity (Fig 2A, B). In contrast, PMA stimulation of *nox2*^*-/-*^ myeloid cells induced no significant ROS activity (Fig 2A-C). This demonstrates that the *nox2* mis-splicing mutation abrogates ROS activity in *nox2*^*-/-*^ mutant myeloid cells.

**Figure 2.**
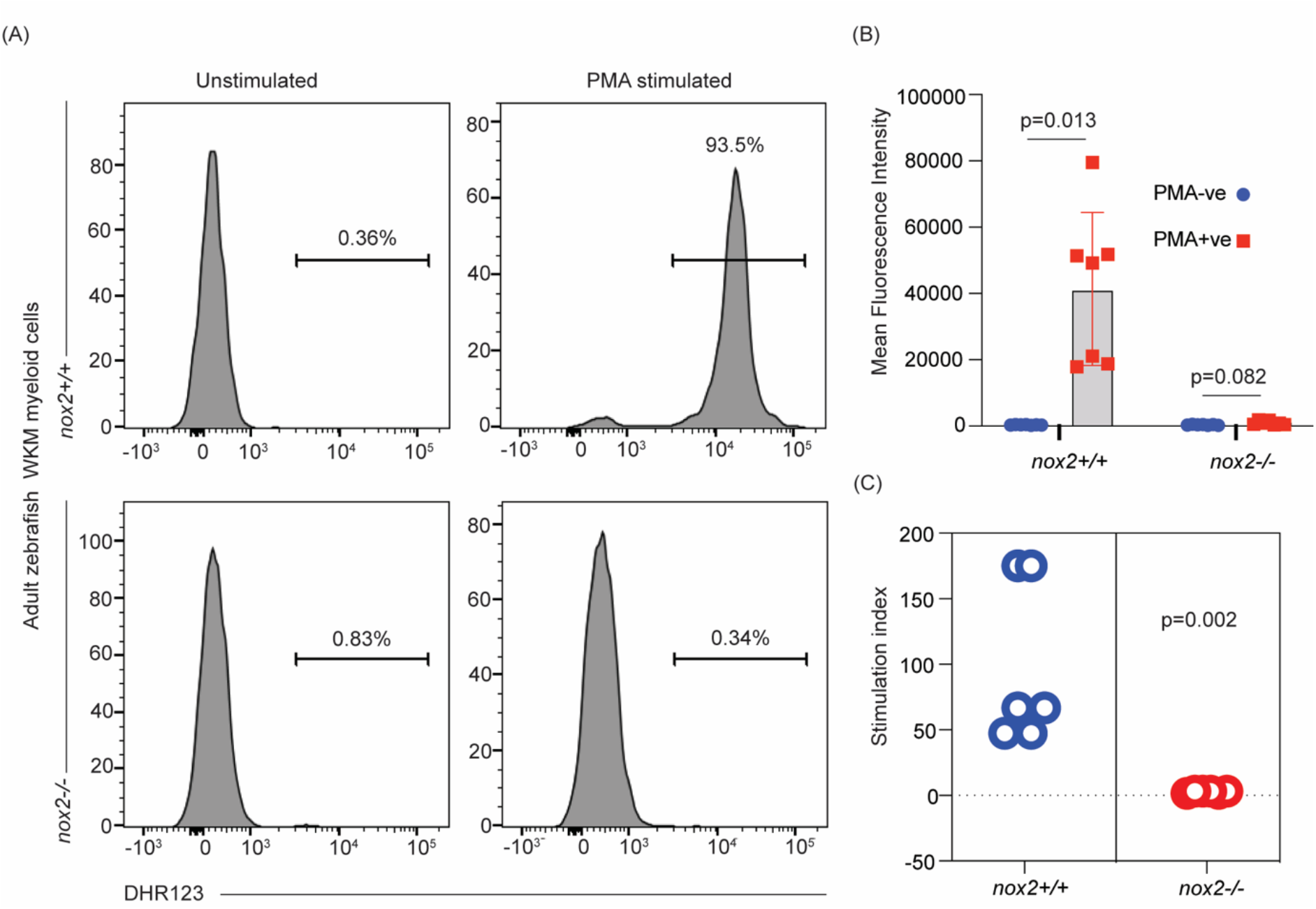
ROS deficiency in adult *nox2-/-* zebrafish myeloid cells. (A)Representative histograms of DHR123 staining flow analysis in *nox2*+/+ (upper panels) and *nox2*-/- (lower panels) adult WKM myeloid cells (B)Comparison of MFI of DHR123 between unstimulated and PMA-stimulated myeloid cells from *nox2*+/+ and *nox2*-/- WKM (C)PMA stimulation index of ROS for *nox2*+/+ and *nox2*-/- WKM myeloid cells DHR123, dihydro rhodamine 123; MFI, mean fluorescence intensity; WKM, whole kidney marrow; +/+, wild type; -/-, homozygous mutant; N, *nox2+/+*, 7; *nox2-/-*, 7. Turkey’s multiple comparison (B); Mann-Whitney (C); p<0.05.

### *nox2* deficiency affects early embryo viability

While mating the *nox2* mutant, a lower-than-expected recovery of embryos carrying the *nox2* mutation for experiments suggested a high rate of early embryonic death. The effect of the *nox2* mutation on early embryo viability was therefore assessed in 5 day survival assays. Concurrent cohorts of wild type, heterozygous and homozygous *nox2* embryos were generated by appropriate crosses: wild type to wild type (WT x WT), wild type to homozygous (WT x *nox2*^-/-^) and homozygous to homozygous (*nox2*^-/-^ x *nox2*^*-/-*^) and monitored for viability. The survival of *nox2* homozygous mutants was significantly reduced, with all excess mortality occurring in the first 24 hpf, regardless of parentage (Fig 3A-C). The mechanism by which *nox2* directly affects early embryo survival is an interesting question for future studies.

**Figure 3.**
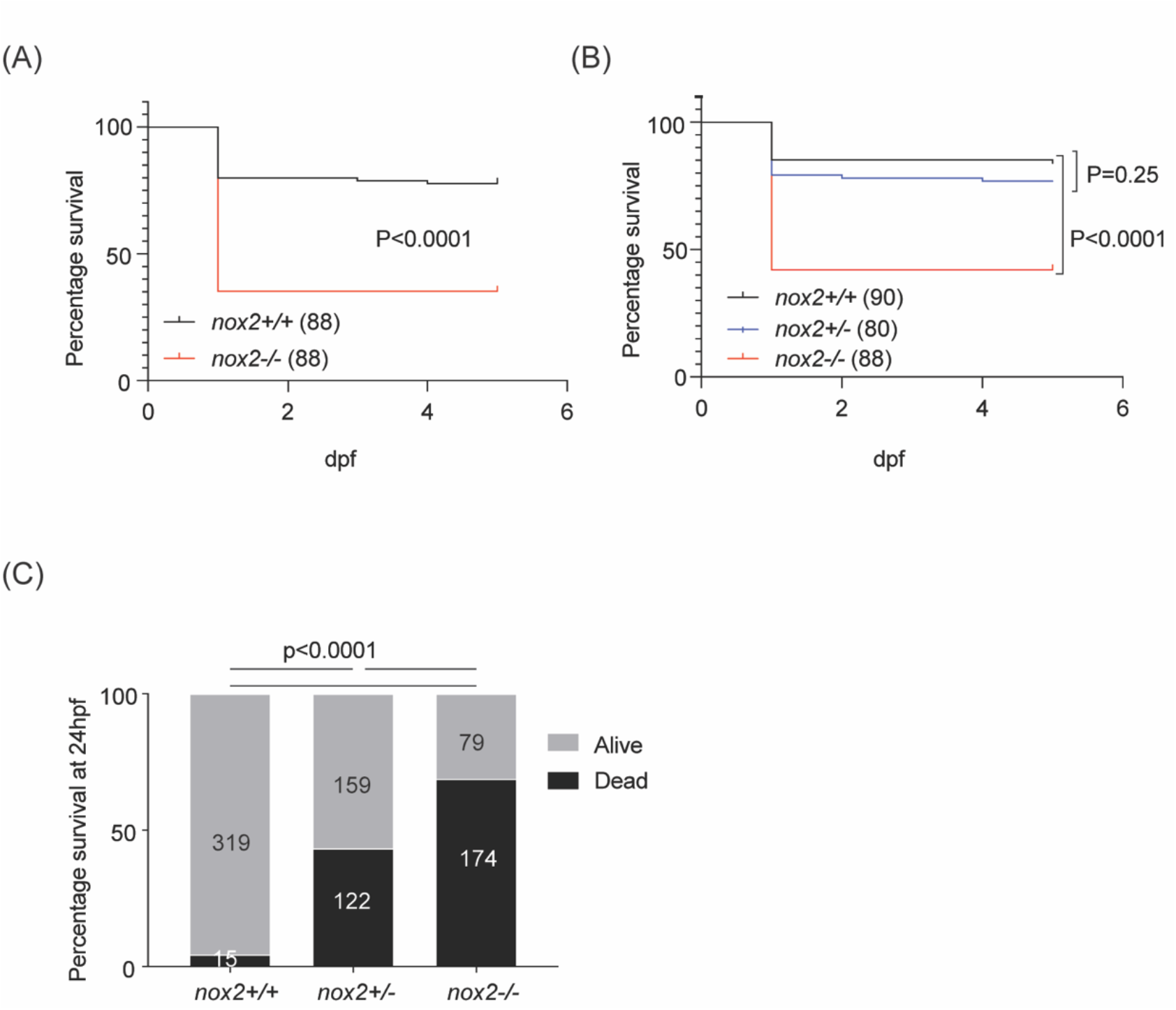
Reduced viability of embryos carrying the *nox2* mutant allele. (A)Survival curve of concurrent cohorts of wild type and *nox2*-/- zebrafish embryos (B)Survival curve of concurrent cohorts of wild type, *nox2*+/- and *nox2*-/- zebrafish embryos (C)Twenty-four (24) hour viability of *nox2* wild type, heterozygous and homozygous zebrafish embryos. A, B and C were performed independently on different days, and represent cohorts of genotype-identical embryos assayed in parallel. N-values in brackets (A, B) and within columns (C) – note that although N-values are similar, A and B are independent experiments; +/+, wild type; +/-, heterozygous; -/-, homozygous. Log-rank (Mantel-Cox) tests (A, B), Fisher’s exact test (C), p <0.0001.

### *nox2* deficiency does not affect initial neutrophil recruitment to a wound

Despite their reduced early viability, enough *nox2* mutant embryos survive for testing biological questions. The arrival of neutrophils at sites of tissue injury is an initial step of acute inflammation, so we investigated the requirement for *nox2* in the initiation of acute inflammation in standard zebrafish injury assays. On the basis of previous reports of NOX2-driven ‘sterile’ hyperinflammation in experimental mice and human patients, we hypothesized that sterile injury would induce excessive neutrophilic inflammation in the *nox2* zebrafish mutants (Dinauer, 2019; Fernandez-Boyanapalli et al., 2010; Zeng et al., 2013).

The temporal profile of neutrophil recruitment to a simple tail fin transection injury is a well-established assay of neutrophil chemotaxis (Renshaw et al., 2006). The *nox2* mutation is carried on transgenic reporter backgrounds allowing neutrophil number and distribution to be easily quantified. The *nox2*^-/-^ embryos had normal and unchanging numbers of trunk neutrophils available for initial relocation to a wound and throughout the 3 hr assay (Fig 4A). Compared to wildtype, there was no consistent statistically significant difference in wound zone neutrophil numbers in *nox2*^+/-^ and *nox2*^*-*/-^ embryos, although a transient statistically significant decrease of uncertain biological significance was observed at 24 hours post wounding (hpw) (Fig 4B).

**Figure 4.**
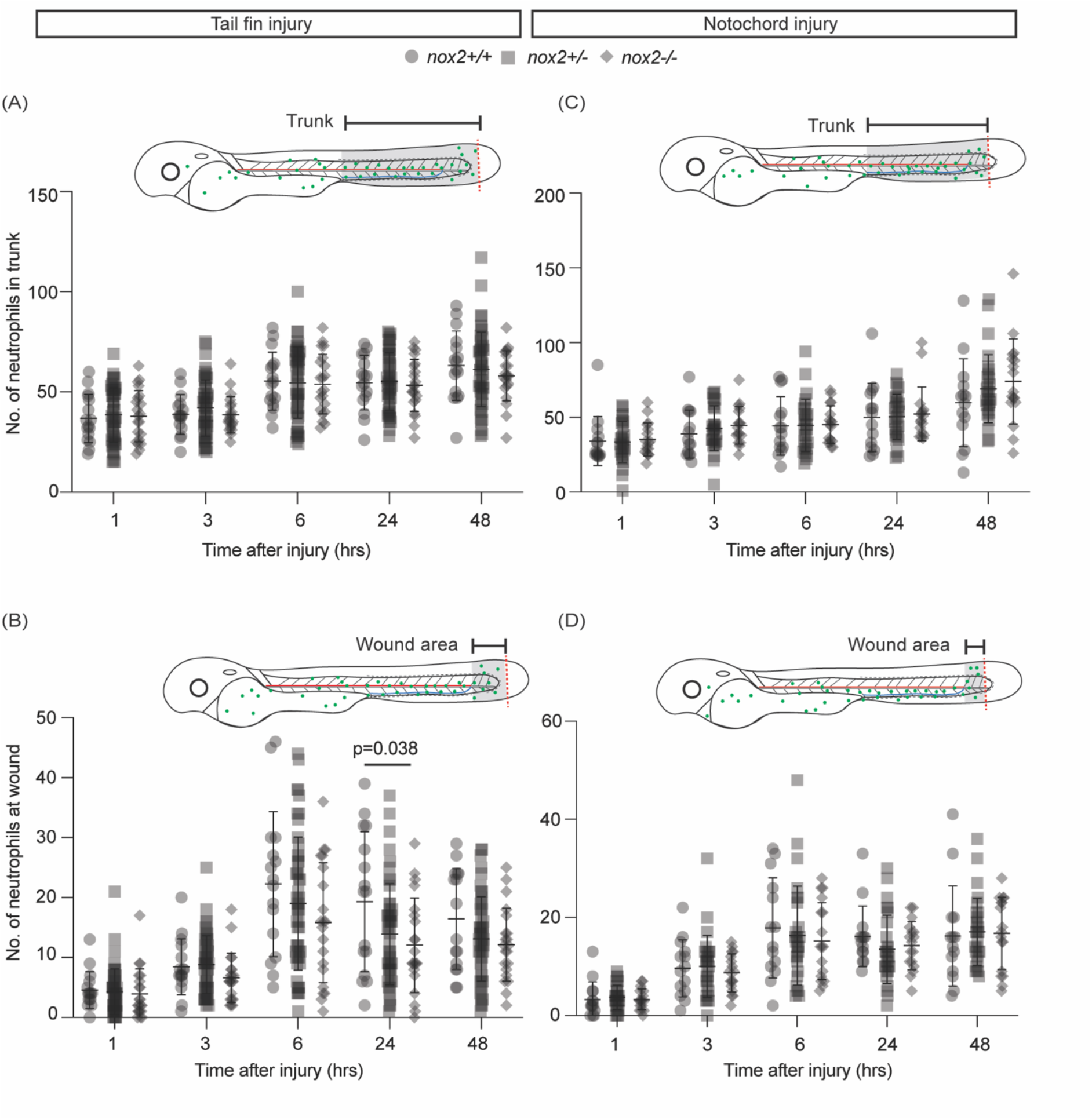
Acute neutrophil inflammatory response in *nox2* zebrafish embryos. (A, C) Number of neutrophils throughout the trunk following tail fin injury (A) and notochord injury (C) (B, D) Number of neutrophils at wound following tail fin injury (A) and notochord injury (D) Tail fin injury, N= 16(+/+), 50(+/-), 21(-/-); notochord injury, N=14 (+/+), 28 (+/-), 17 (- /-); wild type; +/-, heterozygous; -/-, homozygous. Pooled data of three independent experiments. Bars indicate mean ± SD. Two-way ANOVA with Tukey’s multiple comparisons test, p<0.05.

The magnitude of neutrophil migration to tailfin wounds is dependent on the degree of injury, with greater neutrophil inflammatory responses following injuries involving the notochord as well as the fin (Miskolci et al., 2019). We therefore also assessed initial neutrophil recruitment following a larger and more severe tail fin injury involving the tip of the notochord, but again there was no statistically significant difference between wild type and *nox2* mutants (Fig 4 C, D).

These 3-hour assays provided no evidence of augmented initial neutrophil recruitment to an acute sterile injury in *nox2* deficiency states.

## Discussion

We have generated a new *nox2* loss-of-function zebrafish mutant by gene-editing resulting in a mis-spliced *nox2* transcript. Although it remains theoretically possible that this allele, being a splicing mutant, is a hypomorph rather than an absolute null, it is a severely compromising allele and appears to be a functional null. No WT transcript signal was discernable in Sanger sequencing chromatograms of cDNA prepared from *nox2*^*-/-*^ mutants (Fig. 1D). The *nox2*^*-/-*^ mutant neutrophils demonstrated markedly reduced ROS production by neutrophils with a near-zero stimulation index in a standard clinical assay used for CGD diagnosis, functionally validating this allele as a model for the most common form of CGD. *NOX2* mutants account for ∼65% of CGD in human patients (Dinauer, 2019).

There are many *in vitro* studies of NOX2 mutant or NOX-2 functionally-compromised mammalian cells, but only a few animal models (Dinauer, 2019). A murine model replicated the X-linked inheritance of human *NOX2* CGD and displayed the classical CGD disease features of infection vulnerability and an enhanced acute neutrophil response to “sterile” inflammation (Pollock et al., 1995). Two zebrafish *nox2* mutant alleles were reported in a study investigating a *nox2* requirement in retino-tectal development (Weaver et al., 2018). However, despite the truncating frame-shift mutations predicting a severely functionally-compromised protein, neither an *in vivo* biosensor approach or an *in vitro* analysis of cultured retinal ganglion cells were able to detect a significant alteration in hydrogen peroxide dynamics in these mutants in this initial study, although a subsequent study demonstrated a defect (Terzi & Suter, 2020).

However, these two previously-reported zebrafish *nox2* mutant alleles have not been characterized for their effects on myeloid cell ROS production. The functional validation of their ROS production impairment is based on altered H_2_O_2_ dynamics in ex vivo dissociated culture of retinal ganglion cells (Terzi et al., 2021; Weaver et al., 2018). A *p22*^*phox*^*/cyba* zebrafish mutant has been studied for its fungal infection susceptibility; although no direct experimental demonstration of impaired neutrophil ROS production was provided, a neutrophil-specific *p22*^*phox*^*/cyba* rescue reversed a phenotype attributed to ROS deficiency (Schoen et al., 2019). Morphants have also been used to study transient loss of phagocyte NOX subunits, including for example *nox2/cybb* (Bernut et al., 2019; Brothers et al., 2011; Mesureur et al., 2017; Razaghi et al., 2018; Roca & Ramakrishnan, 2013; Yang et al., 2012), *p47*^*phox*^*/ncf1* (Brothers et al., 2013; Brothers et al., 2011; Mesureur et al., 2017; Phan et al., 2018; Sipka et al., 2021; Yang et al., 2012) and *p22*^*phox*^*/cyba* (Prajsnar et al., 2021; Schoen et al., 2019).

The multimeric phagocyte NADPH oxidase enzymatic complex is also of interest as the prototypic NADPH oxidase, and its NOX2 gp91^phox^/CYBB subunit and has provided a paradigm for understanding the structure and function of other NADPH oxidase enzyme complexes (NOX 1, 3-5, and also DUOX 1-2) (Quinn, 2013). Our attempts to validate functional impairment of ROS generation highlight the need to consider both the specificity of the stimulus and the cell types expressing the NADPH oxidase subunit of interest. Although NOX2 is sometimes regarded as a neutrophil-specific PHOX subunit, no individual NADPH oxidase subunit is truly exclusively specific to one cell type (Quinn, 2013). Similarly, while PMA is a potent stimulator of neutrophils (Degroote et al., 2019; Karlsson et al., 2000), in the context of the whole animal it should not be considered to be exclusively a stimulator of ROS produced by the phagocytic oxidative burst (Quinn, 2013).

Hyperinflammation characterized by excessive neutrophil infiltrates and granulomata is a hallmark of CGD (Dinauer, 2019), and is considered to represent a dysfunctional neutrophil inflammatory response in the absence of ROS. However, the mechanisms underpinning this phenomenon are not fully understood. Zebrafish are an ideal model for studying inflammation, as the entire process can readily be visualized in vivo (Robertson & Huttenlocher, 2022), and the acute neutrophil response to stereotypic injuries is well described (Miskolci et al., 2019; Pase et al., 2012; Renshaw et al., 2006). A cell-intrinsic neutrophil requirement for mitochondrial-generated ROS for normal velocity neutrophil migration has been demonstrated (Zhou et al., 2018). Schoen et al (2019) demonstrated an abnormally sustained neutrophil migration response to a fungal infection challenge in a *p22*^*phox*^*/cyba* mutant, using a neutrophil-specific rescue strategy to demonstrate a contribution of a cell-intrinsic neutrophil ROS. Both observations support the hypothesis that there is a cell-autonomous requirement for ROS regulating neutrophil migration. However, we did not observe a quantitative perturbation in the accumulation of neutrophils to two types of “sterile” injury in our *nox2* mutant in the first 3 days after injury, despite the statistical power of pooled group sizes of n>80. It may be that the migratory response needs to be examined over a longer timeframe, or that migration to an ostensibly “sterile” surgical wound involves different mechanisms to those of neutrophil wandering or migration to a fungal infection focus or zymosan-induced “sterile” inflammation (Fernandez-Boyanapalli et al., 2010; Schoen et al., 2019). It may also reflect that the cumulative outcome of the different extents and degrees of ROS production impairment differs in each of these models which are directed at different NADPH oxidase subunits and use different techniques, due to variation in either the full distribution of cell types that have impaired ROS production or in the potential leakiness of the different genetic methodologies. Experiments to explore these interesting possibilities fell beyond the scope of this initial study.

This new *nox2* mutant provides a valuable, well-characterized tool for further exploring this and other *nox2*-dependent phenotypes causing morbidity and mortality for CGD patients.

## Data Availability

Strains are available upon request. The authors affirm that all data necessary for confirming the conclusions of the article are present within the article, figures, and supplementary figures.

## Acknowledgements

We thank Monash AquaCore staff for technical assistance with fish husbandry and Profs. P. Currie for encouragement and S. Renshaw for helpful discussion.

## Contributions

Conceptualization: A.I.I., Z.Z., V.P., G.J.L.; Methodology: A.I.I., Z.Z., V.P.; Formal analysis: A.I.I., Z.Z., G.J.L.; Investigation: A.I.I., Z.Z., Resources: G.J.L.; Data curation: A.I.I., G.J.L; Writing - original draft: A.I.I., G.J.L.; Writing - review & editing: A.I.I., Z.Z., V.P., G.J.L.; Visualization: A.I.I.; Supervision: V.P., G.J.L.; Project administration: G.J.L.; Funding acquisition: G.J.L.

